# The P2Y2 Nucleotide Receptor Mediates Monocyte Tissue Factor Expression and Endotoxemia Death in Mice

**DOI:** 10.1101/2021.12.28.474395

**Authors:** Qianman Peng, Shenqi Qian, Saud Alqahtani, Peter Panizzi, Jianzhong Shen

## Abstract

Recently we reported that in human coronary artery endothelial cells, activation of the P2Y2 receptor (P2Y2R) induces up-regulation of tissue factor (TF), a vital initiator of the coagulation cascade. However, others have shown that monocyte TF is more critical than endothelial TF in provoking a pro-thrombotic state. Thus, we aimed to study whether monocytes express the P2Y2R, its role in controlling TF expression, and its relevance in vivo. RT-PCR and receptor activity assays revealed that among the eight P2Y nucleotide receptors, the P2Y2 subtype was selectively and functionally expressed in human monocytic THP-1 cells and primary monocytes. Stimulation of the cells by ATP or UTP dramatically increased TF protein expression, which was abolished by AR-C118925, a selective P2Y2R antagonist, or by siRNA silencing the P2Y2R. In addition, UTP or ATP treatment induced a rapid accumulation of TF mRNA preceded with an increased TF pre-mRNA, indicating enhanced TF gene transcription. In addition, stimulation of the monocyte P2Y2R significantly activated ERK1/2, JNK, p38, and Akt, along with their downstream transcription factors including c-Jun, c-Fos, and ATF-2, whereas blocking these pathways respectively, all significantly suppressed P2Y2R-mediated TF expression. Furthermore, we found that LPS triggered ATP release and TF expression, the latter of which was suppressed by apyrase or P2Y2R blockage. Importantly, P2Y2R-null mice were more resistant than wild-type mice in response to a lethal dose of LPS, accompanied by much less TF expression in bone marrow cells. These findings demonstrate for the first time that the P2Y2R mediates TF expression in human monocytes through mechanisms involving ERK1/2, JNK, p38, and AKT, and that P2Y2R deletion protects the mice from endotoxemia-induced TF expression and death, highlighting monocyte P2Y2R may be a new drug target for the prevention and/or treatment of relevant thrombotic disease.

## 1. Introduction

Thrombosis is the leading cause of illness and death worldwide. The joint disease caused by acute inflammation and chronic inflammation includes disseminated intravascular coagulation, atherosclerosis, pulmonary embolism, etc. The tissue factor (TF) pathway is essential in secondary hemostasis and is directly involved in the pathophysiology of thrombosis ^1–3^. An abundance of active TF expressed on the dysfunctional endothelial cell surface triggers a hyper-coagulable blood status, resulting in the generation of fibrin and recruitment of activated platelets and neutrophils ^3,4^. Also, diverse stimuli can induce TF expression in monocytes/macrophages, such as C-reactive protein, CD40 ligand, oxidized LDL, PDGF-BB, and endotoxin LPS ^3,5^. TF expression in various cells includes vascular endothelial cells and monocytes in different diseases. Moreover, the exposure of cell surface TF to plasma proteins leads to the binding of factor VIIa to TF, to initiate the process of coagulation, inducing thrombin generation and thrombi formation ^2,3,6,7^. Therefore, identifying novel factors controlling inducible TF expression on blood cells might provide new insights into preventing or treating pathological thrombosis formation.

Over the past few decades, findings indicated that extracellular nucleotides serve as signaling molecules and play an essential role in regulating the cardiovascular system ^8–11^. Extracellular nucleotides generally activate P2Y receptors in inflammatory signaling, although ATP can additionally activate P2X ion channels as well. Eight P2Y G protein-coupled receptor subtypes (P2Y1, 2, 4, 6, 11–14) are identified and well established ^8,12^. Non-platelet P2Y2, P2Y6, and P2Y14 receptors are documented to mediate the inflammatory response in various cell and animal models ^13–18^. Notably, the studies of G protein-coupled P2Y nucleotide receptors produced the clinical anti-platelet drug Plavix targeting the platelet P2Y12 receptor. Except for the extensive research of the P2Y12 receptor in platelets, very little is known about the other seven non-platelet P2Y receptors in inflammation-induced thrombogenesis. Recently we reported that activation of the P2Y2 receptor (P2Y2R) induces up-regulation of TF in human coronary artery endothelial cells ^19,20^. However, others show that the role of monocyte TF is also essential in provoking a pro-thrombotic state ^6,7^. Thus, we propose that the P2Y2R may also mediate TF expression in human monocytes through new mechanisms, highlighting that monocyte P2Y2R could be a new drug target for preventing or treating relevant thrombotic diseases.

Since we have reported that activation of the P2Y2R induces upregulation of TF in vascular endothelial cells ^19,20^, it brings us the interest that monocyte P2Y receptors may also be involved in regulating inducible TF expression. Although the intracellular signaling mechanisms responsible for TF induction are not entirely elucidated, evidence shows that most of these known mediators share similar signal transduction pathways such as MAPK, Akt pathways, and Ca^2+^ signaling ^19–21^. Activation of P2Y receptors is commonly associated with stimulating these pathways, further raising the likelihood of P2Y receptor involvement in monocyte TF induction. Therefore, we hypothesized that activation of P2Y receptor(s) prompts monocyte TF upregulation, leading to the activated monocytes phenotype. The principal objective of this study was to determine whether activation of one or more of the P2Y receptors can induce TF expression in human monocytes and, if so, which P2Y receptor(s) is responsible for this induction. Second, we tried to determine the intracellular signaling pathways involved in P2Y receptor(s)-mediated monocyte TF upregulation and its in vivo relevance. Our findings demonstrate that in human monocytic THP-1 cells and primary monocytes, the P2Y2R is responsible for nucleotides-induced TF expression through multiple signaling pathways, including ERK1/2, JNK, p38, and Akt. In addition, lack of P2Y2R inhibits LPS-induced monocyte TF expression and endotoxemia death in mice.

## 2. Materials and Methods

### 2.1 Materials

Dulbecco’s Modified Eagle Medium (DMEM) and RPMI 1640 were purchased from Lonza. FluoForte™ Kit was purchased from Enzo Life Sciences. Fetal Bovine Serum (FBS) was purchased from Thermo Fisher Scientific. DNA primers were purchased from Integrated DNA Technologies. DNase, RNeasy, and DNeasy kits were purchased from Qiagen. The cDNA synthesis kit was purchased from Applied Biosystems. Purified UTP and ATP were obtained from MilliporeSigma. THP-1 cells were purchased from ATCC. Wild-type and human P2Y2R-transfected 1321N1 cell lines were kindly offered by Dr. Gary Weisman (The University of Missouri-Columbia). Human buffy coats were purchased from BioIVT. Leucosep™ tube was obtained from Greiner bio-one, separation medium, and EasySep human monocytes isolation kits were obtained from Stemcell. All primary and secondary antibodies were obtained from Cell Signaling Technology. siRNA, ON-TARGET plus SMART pool L-003688-00-0005, human P2RY2, NM_002564, and Dharma FECT-1 transfection reagents were purchased from Dharmacon. U0126 and LY294002 were purchased from Cell Signaling Technology. Other reagents were obtained from Tocris.

### 2.2 Cell cultures

#### Human monocytic THP-1 cells

THP-1 cells were cultured in RPMI 1640 medium supplemented with 10% FBS at 37°C in a humidified atmosphere of 5% CO_2_. Before stimulation, cells were seeded to grow for 24 h and starved overnight. Where inhibitor or antagonist was used, cells were pretreated with the inhibitor/antagonist for 45 minutes before cell stimulation.

#### Human primary monocytes

We isolated human primary monocytes from buffy coats using Leucosep™ and a magnetic negative selection kit and seeded the isolated monocytes into the six-well plates in RPMI 1640 medium.

#### Human P2Y2R-transfected 1321N1 cells

Wild-type and human P2Y2R-transfected 1321N1 cells were maintained in DMEM supplemented with 10% FBS at 37°C in a humidified atmosphere of 5% CO_2_. All experiments on transfected cells are run in parallel with wild-type controls.

### 2.3 Intracellular Ca^2+^ mobilization assay

Cells were seeded at a density of 4×10^4^ per well into 96-well culture plates and cultured for one day. On day two, the original medium was removed. The assay medium from the FluoForte™ kit containing the Ca^2+^ dye was added, and receptor-mediated Ca^2+^ mobilization was determined as previously described ^19,22^. Fluorescence was determined immediately after adding different reagents, with an excitation wavelength set to 490 nm and an emission wavelength set to 525 nm, and readings were taken every 1s for 640s. For the antagonist inhibition experiment, cells were pre-incubated with the antagonist for 45 min before agonist addition. Measurement of Ca^2+^ signal was performed with the fluorometer plate reader (BMG FLUOstar), the results of which were shown as relative fluorescence units (RFU).

### 2.4 ATP release assay

ATP release from THP-1 cells was determined by the ENLIGHTEN ATP Assay System Bioluminescence Detection Kit (Promega). Cells were grown in 6-well plates and starved overnight. Before performing the ATP assay, a fresh basal culture medium was added with the NTPDase inhibitor ARL67156 obtained from Tocris (100μM, 30min) into each well. Then, the accumulation of released ATP was stimulated by LPS (100ng/ml) treatment for the indicated time. 100μL medium from each well of the 6-well plate was transferred to an opaque-wall 96-well plate. Next, use a multiple channel pipette to add 100μL assay reagent into each well, following the kit’s protocol. Measurement of ATP concentration was performed with the Glomax 96 microplate luminometer (Promega), the results of which were shown as relative light units (RLU).

### 2.5 RT-PCR and Real-time PCR analysis

Total RNA and DNA were extracted from THP-1 cells or primary human monocytes using the RNeasy and DNeasy kits, respectively. On-column DNA digestion was carried out during RNA extraction. For the synthesis of the first strand of cDNA, 1 μg of total RNA after DNase treatment was reverse transcribed using a cDNA synthesis kit. The cDNA samples were then amplified by PCR using 2.5 units of Taq DNA polymerase (Qiagen). Real-time PCR was performed on a CFX96 detection system (Bio-Rad) with SYBR Green reagents (Qiagen). All PCR primers’ sequences are presented in online supplemental table 1.

### 2.6 Western blotting assays

After stimulation, cells were lysed, and standard Western blotting was performed as described previously ^19,20^. The individual primary antibodies used were anti-p-ERK1/2, anti-p-p38, anti-p-SAPK/JNK., anti-p-AKT, anti-p-c-Jun, anti-p-c-Fos, anti-p-ATF-2, and anti-tissue factor. Equal protein loading was verified by stripping off the original antibodies and re-probing the membranes with the primary antibodies anti-ERK1/2, anti-p38, anti-SAPK/JNK, anti-AKT anti-β-tubulin (Cell Signaling Technology).

### 2.7 Silencing of P2Y2R by siRNA

The procedure of knocking down the P2Y2R was performed as described previously ^19^. THP-1 cells were transfected using the four-sequence siRNA pool (ON-TARGET plus SMART pool L-003688-00-0005, human P2RY2, NM_002564, Dharmacon) and the Dharma FECT-1 transfection reagent following the manufacturer’s protocol. Briefly, THP-1 cells were seeded in 6-well plates at 80–90% confluence; the medium was replaced with complete RPMI 1640 without antibiotics before transfection. DharmaFECT-1 transfection reagent and siRNA were incubated separately at room temperature for 5 min. Mixtures were combined, incubated another 20 min, and added to cells at a final concentration of 5μl/ml DharmaFECT 1 and 50 nM P2Y2R-targeting or scrambled siRNAs. Real-time PCR assays were performed to confirm the decrease of P2Y2R mRNA after 24 h post-transfection.

### 2.8 Endotoxemia mouse model and survival study

Wild-type (C57BL/6) and P2Y2R mutant (P2Y2R^−/−^) on a C57BL/6 background were purchased from The Jackson Laboratory (Bar Harbor) and bred at the animal facilities at Auburn University. DNA extraction from mouse tail and PCR genotyping were routinely performed. Mouse P2Y2R wild-type primer: 5’-AGC CAC CCG GCG GGC ATA AC-3’; Mouse P2Y2R mutant: 5’-AAA TGC CTG CTC TTT ACT GAA GG-3’; Mouse P2Y2R common primer: 5’-GAG GGG GAC GAA CTG GGA TAC-3’. Male C57BL/6 mice in each group (2-3 months old, 25-35g) were challenged with LPS intravenously via tail vein. Injections were made in P2Y2R^+/+^ and P2Y2R^−/−^ mice groups with 100 µl of saline containing 900 µg of LPS. The data represent the number of animals surviving within five days after LPS injections. All mice were maintained in a facility free of well-defined pathogens under the supervision of the Animal Health Unit at Auburn University, College of Veterinary Medicine. All animal protocols were approved by the Institutional Animal Care and Use Committee at Auburn University.

### 2.9 Isolation of mouse bone marrow cells

Mouse bone marrow cells were isolated from mice 24 h after LPS injection. Cell debris was removed by centrifugation, and the cell lysates were prepared by homogenization (BeadBug™ mini homogenizer model D1030, Benchmark) in Laemmli Lysis Buffer (Sigma). Protein concentration was measured by Nanodrop 2000c (Thermo).

### 2.10 Data analysis

All data were analyzed by Prism 9 (GraphPad Software Inc). Data are expressed as the mean ± SEM. The means of two groups were compared using Student’s t-test (unpaired, two-tailed), and one-way analysis of variance was used for comparison of more than two groups with P < 0.05 considered to be statistically significant. Unless otherwise indicated, all experiments were repeated at least three times.

## 3. Results

### P2Y receptor mRNA expression in THP-1 and primary monocytes

We first analyzed the expression profile of P2Y receptors in human monocytic THP-1 cells and human primary monocytes, as it has not been entirely determined. Our RT-PCR analysis showed that THP-1 cells expressed P2Y1, P2Y2, and P2Y6 receptor mRNAs, with no or barely detectable mRNAs for the other five subtype receptors (Fig. 1A). However, human primary monocytes expressed P2Y2R mRNAs, with virtually no detectable mRNAs for the P2Y4 or P2Y6 receptors (Fig. 1B). This result indicates that human monocytes predominately express the P2Y2R among the P2Y receptor family.

**Figure 1.**
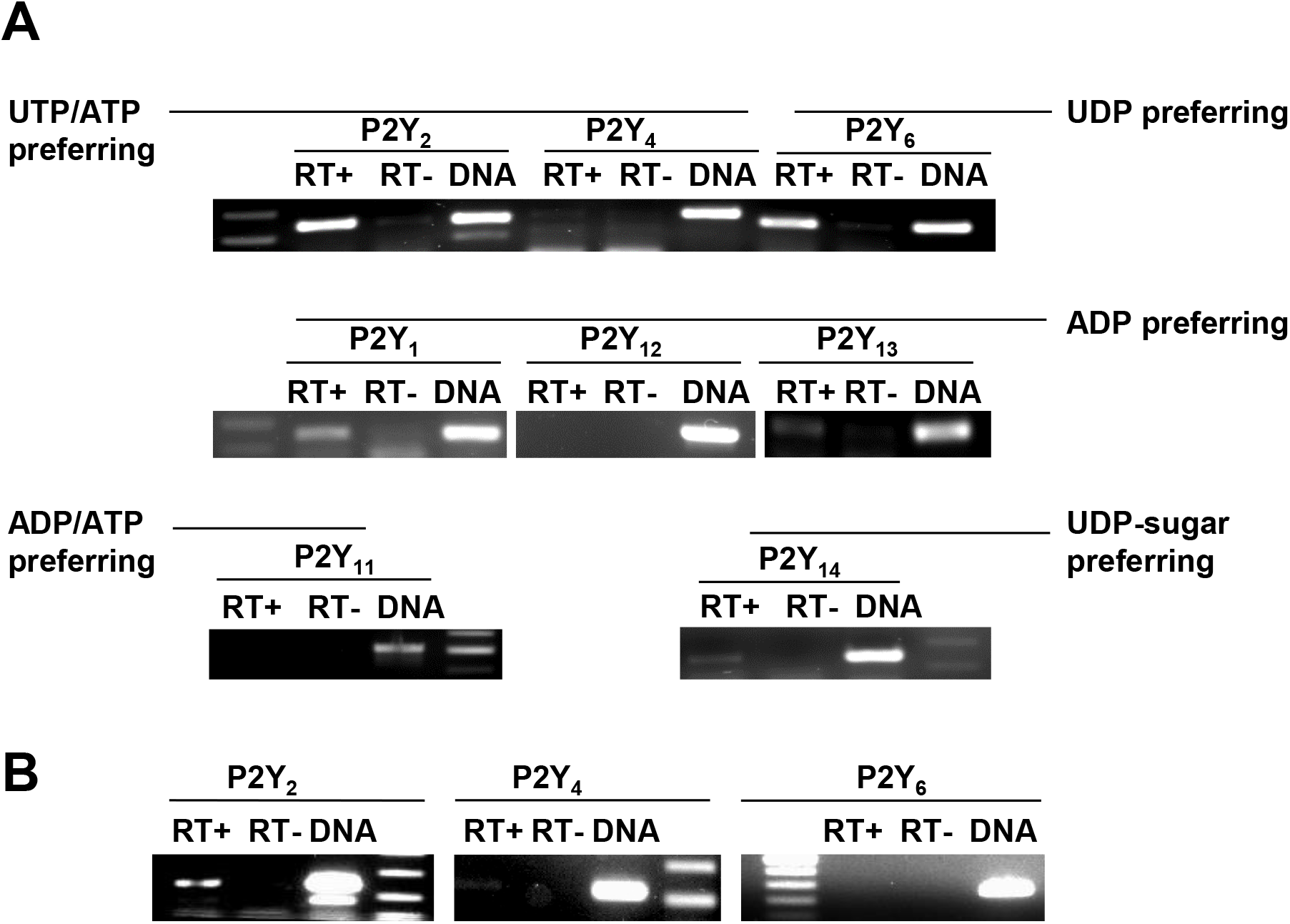
Expression profile of P2Y receptors in human monocytic THP-1 cells and primary monocytes. mRNA expression of eight subtypes of P2Y receptors in THP-1 monocytes (**A**) and human primary monocytes (**B**) was determined by RT-PCR. RT-stands for RT-PCR performed without reverse transcriptase. Genomic DNA was used as the positive control. The data shown represent three independent experiments.

### The P2Y2R mediates nucleotides-induced Ca^2+^ signaling in THP-1 cells

To differential the role of P2Y2R, P2Y1R, and P2Y6R in post-receptor Ca^2+^ signaling in THP-1 cells, we pretreated cells with P2Y2R agonist UTP and ATP, P2Y1R agonist ADP and 2-Me-ADP, and P2Y6R agonist UDP and MRS 2693 (Fig. 2A). UTP and ATP 10μM triggered a peak Ca^2+^ response. 10μM ADP or 10μM UDP activated peak Ca^2+^ responses, but they showed a much weaker response. However, the P2Y1R agonist, an ADP analog, 2-Me-ADP did not induce significant Ca^2+^ mobilization even at 100μM, and the P2Y6R-selective agonist MRS-2693 at 100 μM did not induce any Ca^2+^ response. To test whether ADP and UDP can induce P2Y2R-mediated Ca^2+^ mobilization, we tested UTP, ATP, UDP, and ADP at 10μM on P2Y2R-transfected 1321N1 cells and wild-type cells lacking any P2Y and P2X receptors (Fig. 2B). Interestingly, UTP, ATP, UDP, and ADP significantly induced Ca^2+^ mobilization in P2Y2R-transfected 1321N1 cells, but UDP/ADP treatment showed less effectiveness than UTP/ATP did (Fig. 2B). To confirm that the tested nucleotides selectively activate the P2Y2R, we pretreated wild-type 1321N1 cells with the four nucleotides. Fig. 2B showed that UTP, UDP, ATP, and ADP all had no effect on Ca^2+^ mobilization in wild-type 1321N1 cells, but Carbachol showed a positive response as a control. To further determine whether UDP- and ADP-induced Ca^2+^ mobilization is mediated by P2Y2R in THP-1 cells, we pretreated THP-1 cells with 3 μM ARC-118925XX, a P2Y2R-selective antagonist. As expected, ARC-118925 suppressed intracellular Ca^2+^ elevation caused by UDP and ADP (Fig. 2C). In contrast, pretreatment of the THP-1 cells with a specific inhibitor of P2Y1R MRS-2179 (1μM) or with a specific inhibitor of P2Y6R MRS-2578 (1μM) did not affect ADP- and UDP-induced peak Ca^2+^ response in THP-1 cells (Fig. 2D). These data indicate that the P2Y2R mediated ADP-, UDP-, ATP-, and UTP-induced Ca^2+^ signaling in THP-1 cells.

**Figure 2.**
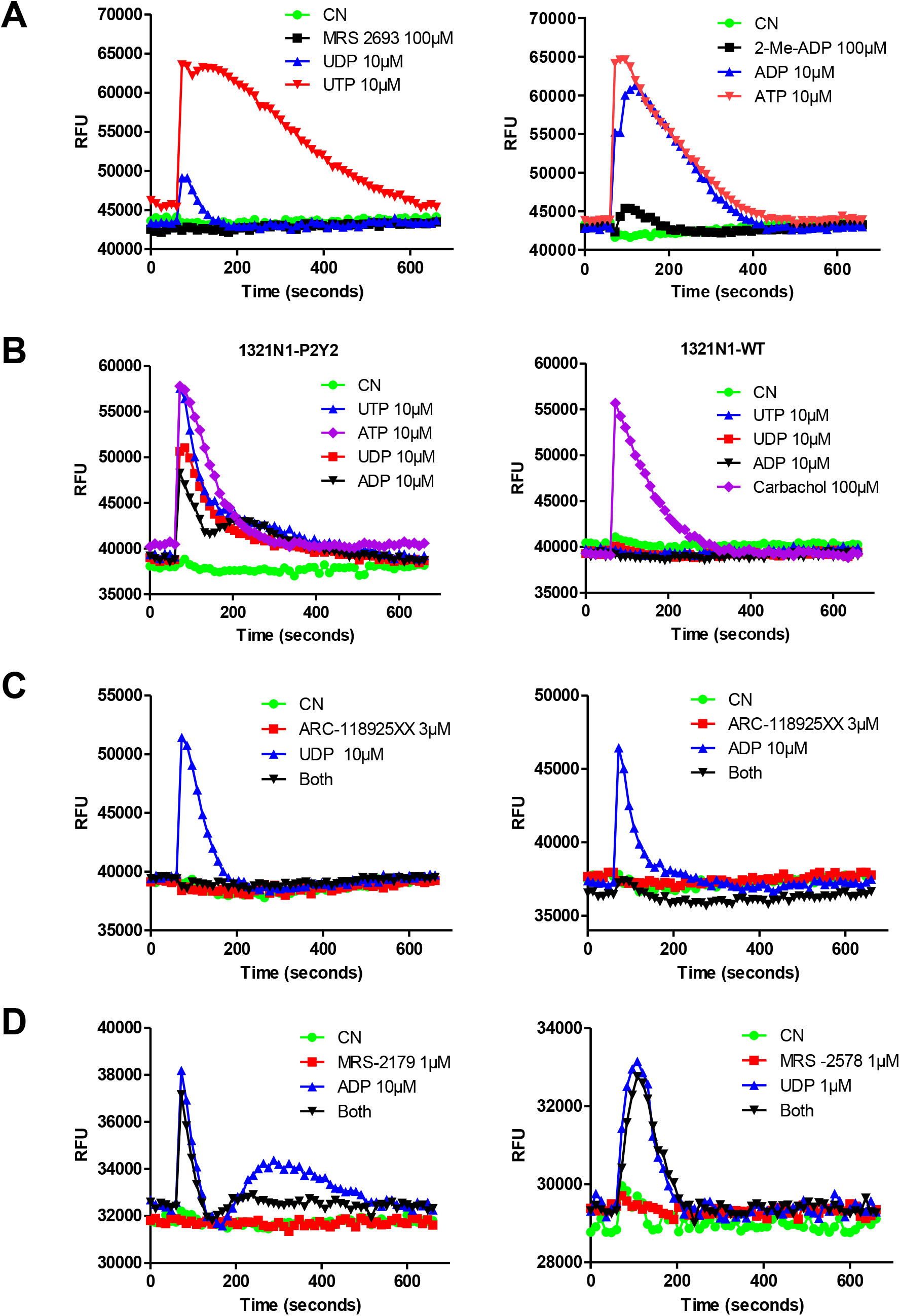
P2Y2R mediation of nucleotides-induced Ca^2+^ signaling in THP-1 cells. P2Y1 receptor agonist ADP- and 2-Methylthio-ADP (2-Me-ADP)-, P2Y6 receptor agonist UDP-and MRS-2693-induced intracellular [Ca^2+^] mobilization were determined in THP-1 cells, UTP and ATP were used for controls (**A**). ATP-, UTP-, ADP-, and UDP-induced intracellular [Ca^2+^] mobilization were determined in 1321N1 wild-type cells and cells transfected with P2Y2R; Carbachol was used as a positive control (**B**). Effect of P2Y2R-selective antagonist ARC-118925XX on ADP- and UDP-induced [Ca^2+^] mobilization in THP-1 cells (**C**). Effect of P2Y1R-selective antagonist MRS-2179 on ADP-induced intracellular [Ca^2+^] mobilization and effect of P2Y6R-selective antagonist MRS-2578 on UDP-induced intracellular [Ca^2+^] mobilization (**D**). Changes in Ca^2+^ signal are shown as relative fluorescence units (RFU; y-axis). The data shown are representative of three independent experiments performed in triplicate wells of each condition.

### Extracellular nucleotides-induced TF upregulation

To determine whether activation of the P2Y2R can induce TF expression, THP-1 cells were stimulated with agonists for various time intervals. TF protein expression was increased within 1 h, reached a maximum level at 4 h, and returned to the basal level at 8 h after the serum-starved cells were challenged with 100 μM ATP or UTP (Fig. 3A). A dose-dependent increase of TF protein expression was observed after the cells were stimulated with different concentrations of ATP or UTP (Fig. 3B). Of note, the level of TF induction by UTP/ATP was even comparable to that of 100ng/ml LPS (<u>Fig</u>. 3B), a known inducer of TF expression in various cells. Similarly, TF protein expression was increased in human primary monocytes with 100 μM ATP or UTP stimulation (Fig. 3C). In addition, stimulation of the cells by ADP or UDP increased TF expression as well, which was suppressed by ARC-118925 pretreatment (Supplemental figure 1).

**Figure 3.**
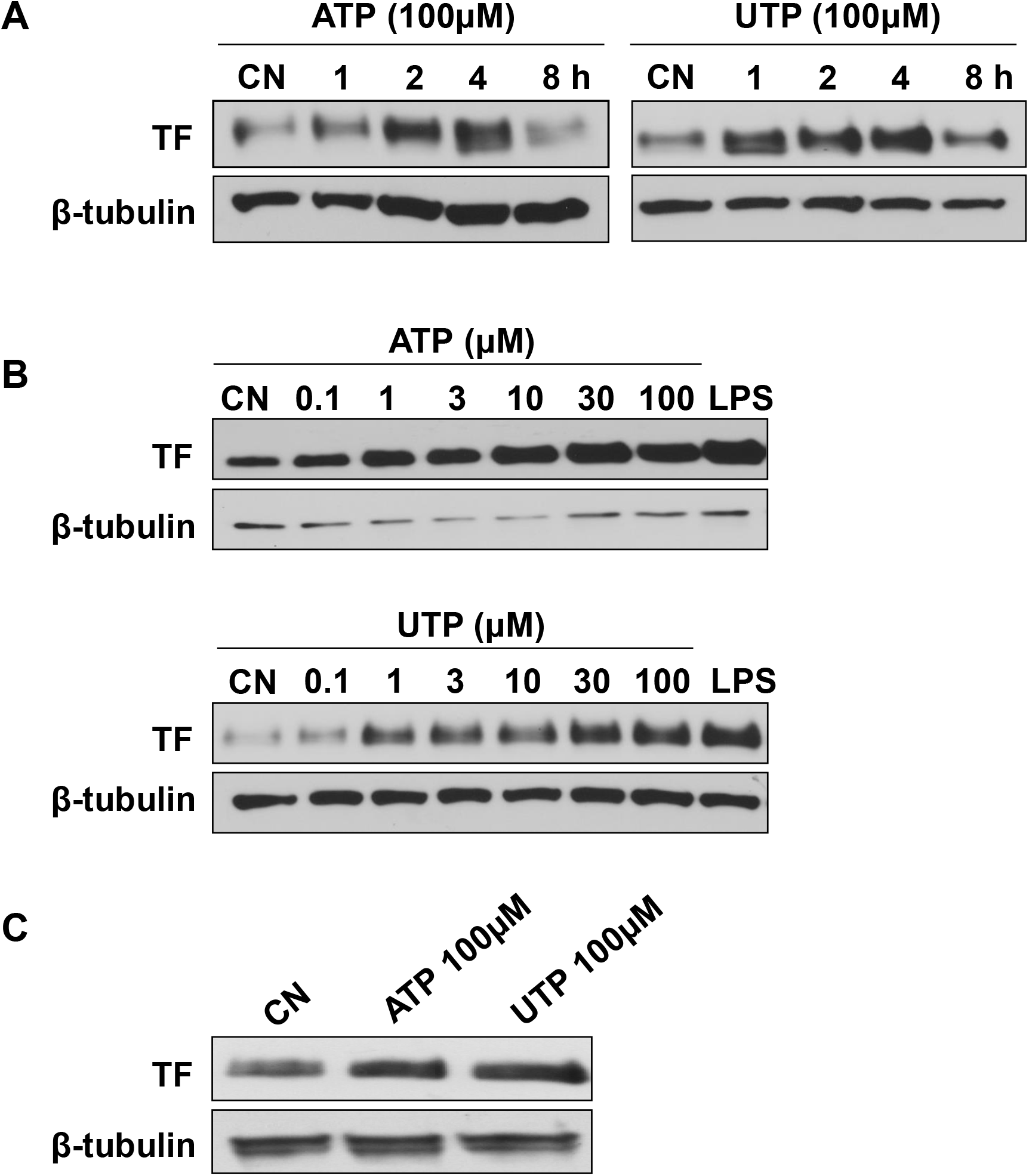
Extracellular UTP/ATP upregulation of TF expression. Total cellular TF protein expression was determined by Western blotting assay in THP-1 cells stimulated by 100 μM UTP or ATP for the indicated times (**A**) or by the stated UTP/ATP concentrations for 4 hours (**B**). Total cellular TF protein expression was determined by Western blotting assay in human primary monocytes stimulated by 100 μM UTP or ATP for 4 hours (**C**). LPS was used as a positive control. The data shown represent three independent experiments.

### UTP/ATP up-regulation of TF mRNA expression

Next, we performed real-time PCR assays to determine whether TF pre-mRNA and mature mRNA are affected by UTP/ATP stimulation. An elevation of TF pre-mRNA level was detected as early as 1 h, and expression maintained 2-fold of the basal level after 4 h. A peak elevation of TF mature mRNA level was detected at 2 h for 100 μM UTP and 3 h for ATP (Fig. 4A). TF mRNA level was also upregulated dose-dependent by ATP or UTP (Fig. 4B). This result indicates that UTP/ATP-sensitive P2Y2R activation induces TF mRNA transcription.

**Figure 4.**
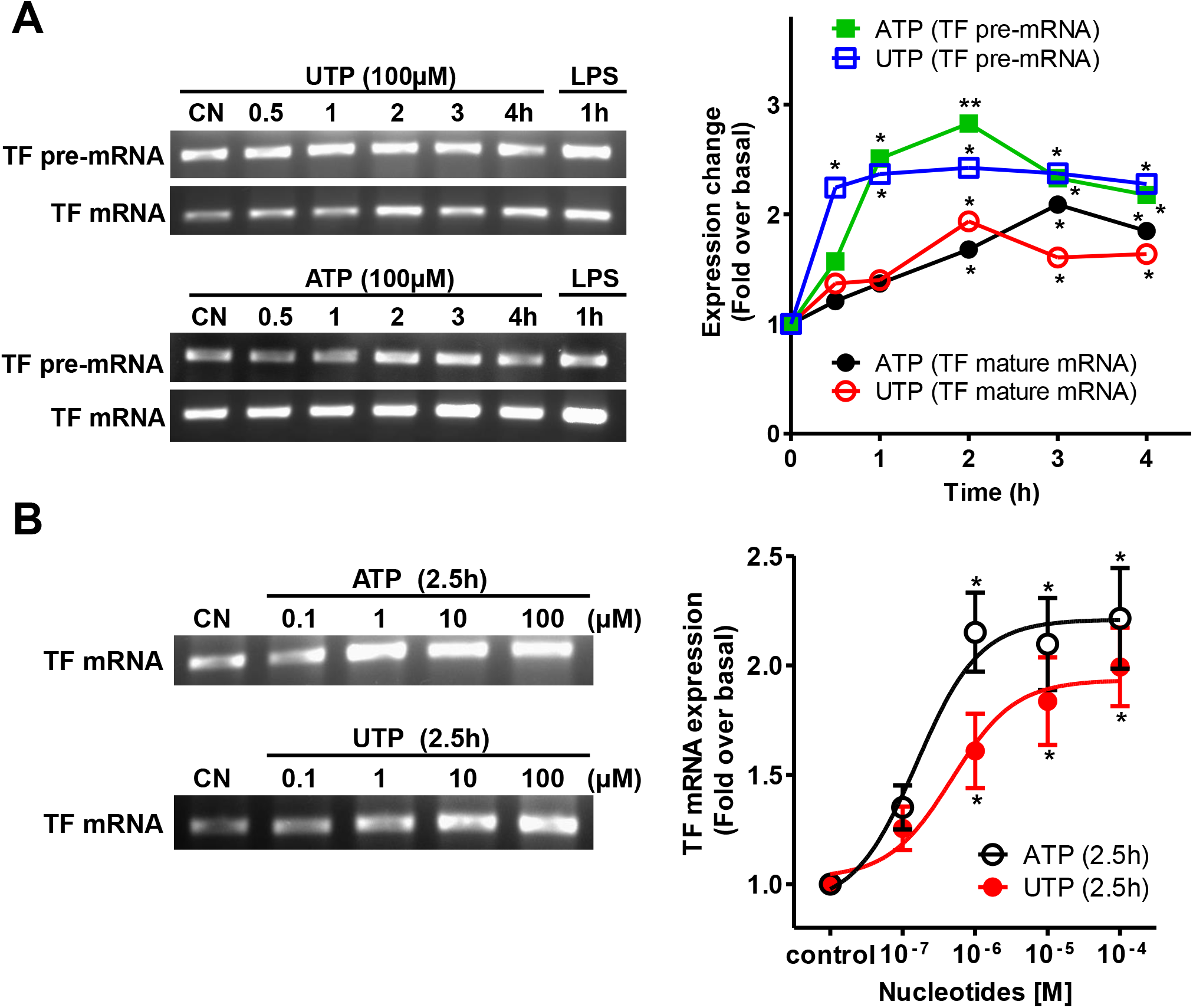
P2Y2R activation by UTP/ATP promotes TF pre- and mature mRNA expression. Two different pairs of primers were designed to quantify human TF mature mRNA and pre-mRNA levels by Real-time PCR assays. Representative RT-PCR gel images show the time-dependent effect of UTP and ATP on TF pre-mRNA and mature mRNA expression, LPS 1h treatment as a positive control, and GAPDH mRNA as an internal control (not shown). Data were summarized from three independent experiments. *, p < 0.05; **, p < 0.01, relative to the 0 time point (**A**). TF mature mRNA was analyzed by real-time PCR after THP-1 cells were stimulated with 100 μM ATP or UTP for 2.5h. Representative RT-PCR gel images show the dose-dependent effect of UTP and ATP on the expression of TF mature mRNA. GAPDH was used as a control (not shown). Data were summarized from three independent experiments. *, p < 0.05, relative to the unstimulated vehicle control (**B**).

### The P2Y2R mediates UTP/ATP-induced TF expression

To determine whether the nucleotides’ action is through P2Y2R, ARC-118925XX, a selective antagonist of P2Y2R, was employed. Fig. 5A shows that 3μM ARC-118925XX pretreatment abolished UTP/ATP-induced TF protein expression, suggesting an involvement of the P2Y2R. To further confirm this, we knocked down the P2Y2R by a siRNA approach. Fig. 5B shows that P2Y2R mRNA decreased about 50% after transfection of a pool of P2Y2R siRNAs for 24 h, while the scrambled siRNA had no effect as determined by real-time PCR. Knocking down P2Y2R dramatically inhibited ATP-induced Ca^2+^ signaling compared to scrambled siRNA treatment (Fig. 5C). Surprisingly, silencing P2Y2R abrogated not only UTP-induced but also LPS-induced TF expression compared with scrambled siRNA treatment (Fig. 5D). Collectively, these data indicate that the P2Y2R mediates nucleotides-induced and possibly LPS-induced TF expression as well in THP-1 cells.

**Figure 5.**
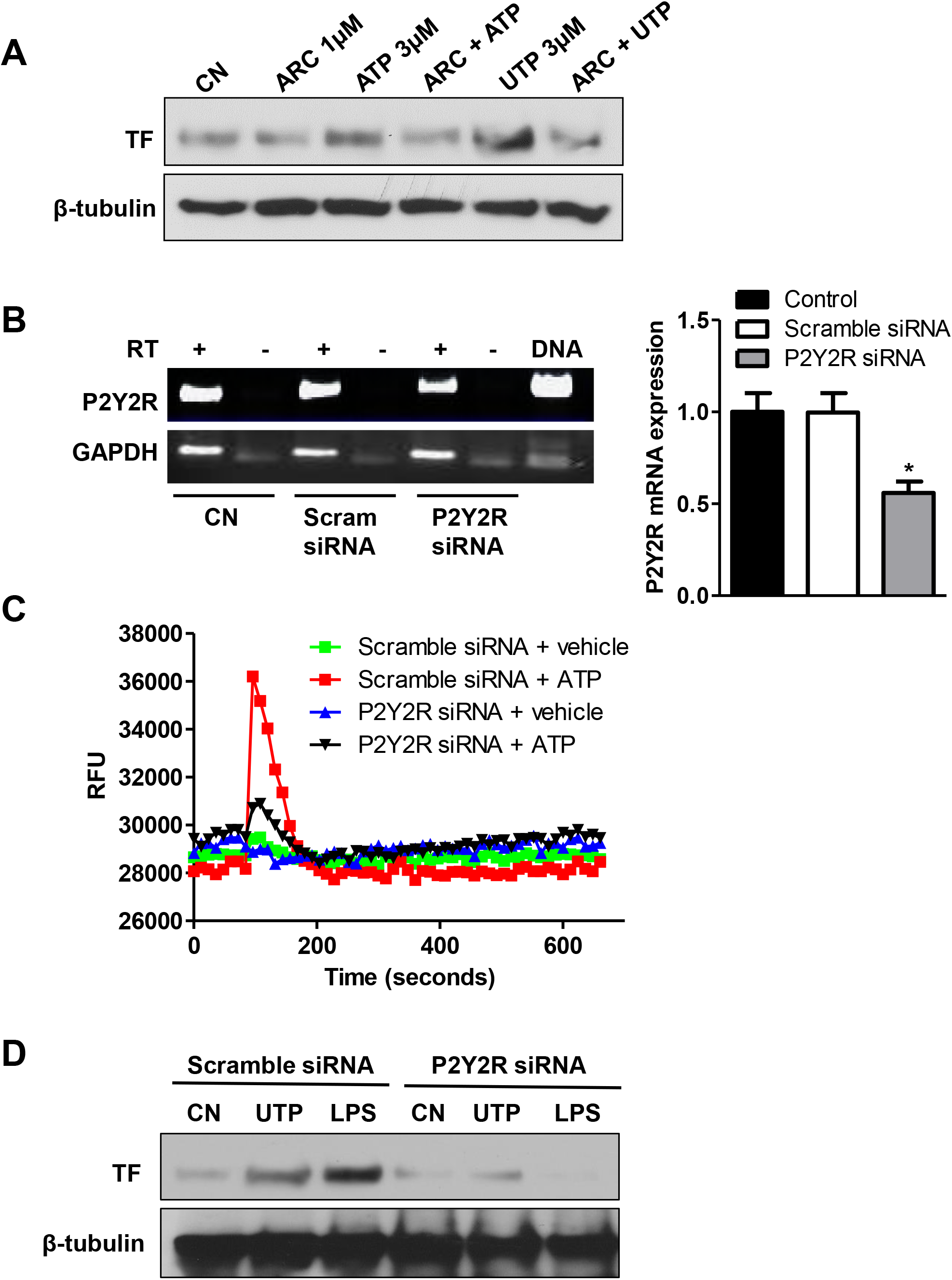
P2Y2R mediates UTP/ATP-induced TF expression. TF protein expressions were analyzed by Western blotting after the cells were pretreated with ARC-118925XX (ARC, 1 μM, 40 min), a P2Y2R-selective antagonist, followed by ATP or UTP stimulation of the cells for 4 h (**A**). P2Y2R mRNA expressions were determined by real-time RT-PCR after THP-1 cells were transfected with P2Y2R-siRNA or scrambled control siRNA for 24 h. Data were summarized from three independent experiments. *, p < 0.05 relative to control (**B**). UTP and ATP (10 μM)-induced intracellular [Ca^2+^] mobilization was determined in THP-1 cells transfected with P2Y2R-siRNA or scrambled control siRNA for 24 h. The data shown are representative of three independent experiments (**C**). The Western blotting assay evaluated UTP-stimulated (4h) TF protein expression after the THP-1 cells were transfected with P2Y2R-siRNA or scrambled control siRNA for 24 h. LPS treatment (100ng/ml for 4h) as a control. The data shown are representative of three independent experiments (**D**).

### Role of the P2Y2R in ATP/UTP-induced MAPK and AKT cell signaling

Next, we studied post-P2Y2R signaling. Fig. 6A and 6B showed that UTP/ATP caused rapid phosphorylations of ERK1/2, p38, JNK, and AKT in a dose-dependent manner. Interestingly, UTP/ATP induced MAPK pathways from 0.1 μM and reached maximal phosphorylation at 1 μM. However, further UTP/ATP concentration increased to 10 μM or above led to less phosphorylation of ERK1/2, p38, and JNK, but not AKT. This might be explained in part by a G protein in reverse mode. High concentrations of agonists might catalyze GTP dissociation from activated heteromeric Gαβγ and lead to a nucleotide-free G protein sequestrated in heterotrimeric conformation at the active P2Y2R thus attenuating downstream signaling in an agonist-dependent manner ^23^. In contrast, ATP/UTP-induced AKT signaling showed a typical dose-dependent fashion. Fig. 6C shows that UTP/ATP-induced MAPK and AKT activation was suppressed by ARC-118925XX, suggesting a role of P2Y2R in UTP/ATP signaling. Moreover, Fig. 6D shows that UTP treatment induced rapid phosphorylations of c-Fos, c-Jun, Fra-1, and ATF-2 in a time-dependent manner.

**Figure 6.**
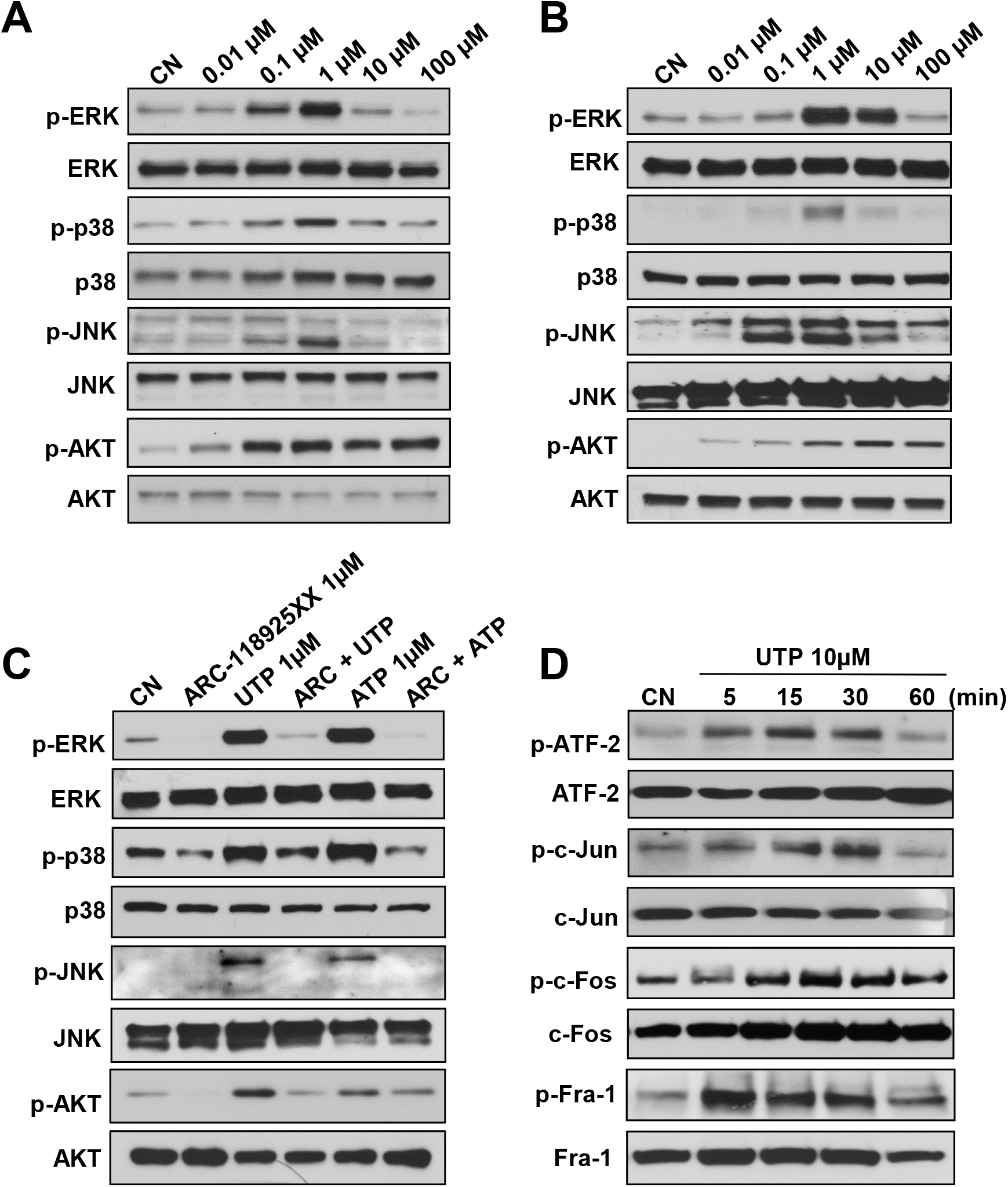
P2Y2R-mediated MAPK and AKT signaling. Phosphorylation levels of ERK1/2, p38, JNK, and AKT were evaluated by Western blotting assay after THP-1 cells were stimulated by indicated concentrations of UTP (**A**) and ATP (**B**) for 10 min. Phosphorylation levels of ERK1/2, p38, JNK, and AKT were analyzed by Western blotting assay after THP-1 cells were pretreated for 45 min with ARC-118925XX followed by UTP or ATP stimulation of the cells for 10 min (**C**). Phosphorylation levels of AP-1 subunits, c-Jun, c-Fos, Fra-1, and ATF-2 in response to P2Y2R activation were detected by Western blotting assay after THP-1 cells were stimulated by 10 μM UTP for the indicated times. The data shown are representative of three independent experiments (**D**).

### Role of MAPK and AKT signaling pathways in P2Y2R-mediated TF expression

We then further explored the contribution of the MAPK and AKT signaling pathways on TF induction. Fig.7 shows that UTP/ATP-induced phosphorylations of ERK1/2, p38, JNK, and AKT were abolished by the MEK1/2 inhibition with the MEK inhibitor U0126 (Fig. 7A), the p38 MAPK inhibitor SB203580 (Fig. 7B), the JNK kinase inhibitor SP600125 (Fig. 7C), and the PI3 kinase inhibitor LY294002, respectively (Fig. 7D). In addition, pretreatment of the cells with the individual four inhibitors all significantly suppressed UTP-or ATP-induced TF upregulation (Fig. 7A to 7D).

**Figure 7.**
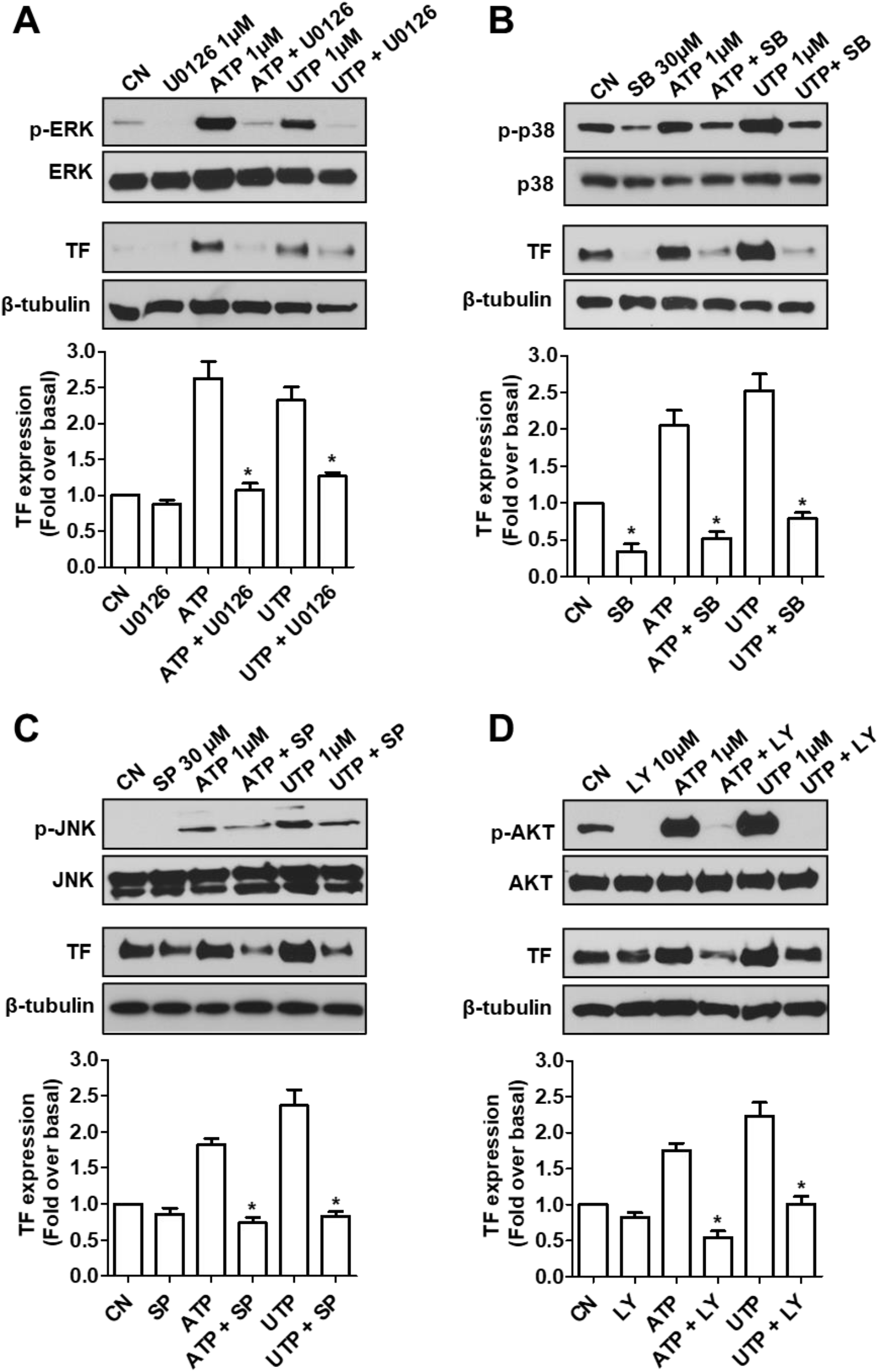
Effect of MAPK and PI3K/AKT pathways on P2Y2R-induced TF expression. TF protein expressions were determined by Western blotting assays after the THP-1 cells were pretreated with the individual inhibitor for ERK1/2 (U0126, **A**), p38 (SB203580, **B**), JNK (SP600125, **C**), or PI3K (LY294002, **D**) for 45 min followed by ATP or UTP stimulation for 4 h. TF data were summarized from three independent experiments. *, p < 0.05, relative to the respective controls. Data on the individual inhibitor’s effective blockage of the MAPK and AKT pathways were presented at the top of each panel.

### Role of the P2Y2R in LPS-induced TF upregulation and endotoxemia death in mice

Finally, we explored the disease relevance of the proposed P2Y2R/TF axis. First, we found that LPS treatment dramatically induced ATP release from 1 to 2 h in THP-1 cells (Fig. 8A). Second, we discovered that removing extracellular nucleotides by apyrase inhibited LPS-induced TF induction (Fig. 8B). Third, blocking the P2Y2R by two structurally different P2Y2R antagonists (ARC-118925XX and Kaempferol) similarly abolished LPS-induced TF upregulation, suggesting a significant role of the P2Y2R. To further investigate the in vivo role of P2Y2R in endotoxemia, we challenged the wild-type and age-matched P2Y2R-null mice with a lethal dose of LPS. Fig. 8E shows that the P2Y2R-null mice exhibited a significantly higher survival proportion than the wild-type mice in response to the same LPS challenge. We also collected bone marrow cells from both groups of mice, and Western blotting assays showed that the wild-type mouse bone marrows expressed a much higher level of TF in response to LPS challenge in vivo than the bone marrows isolated from P2Y2R-null mice (<u>Fig.</u> 8F).

**Figure 8.**
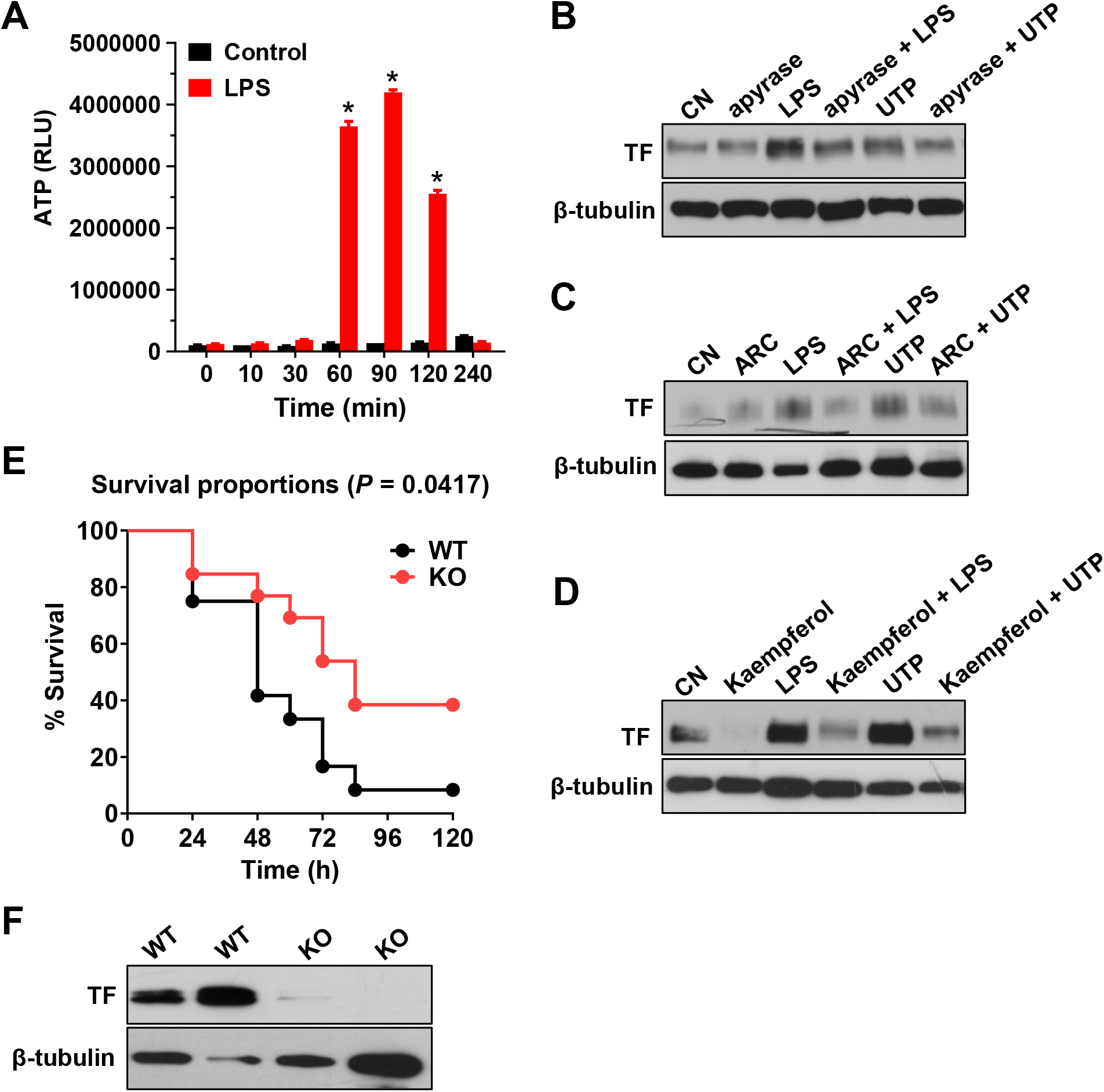
Role of P2Y2R in LPS-induced TF upregulation and endotoxemia death in mice. LPS (100ng/mL)-induced ATP release from cultured THP-1 cells was evaluated for the indicated times (**A**). LPS (100ng/mL)-induced and UTP (10 μM)-induced TF protein (4 h) expressions in THP-1 cells were analyzed by Western blotting assays after the cells were pretreated with apyrase (5 U/ml) for 45min (**B**). LPS (100ng/mL)-induced and UTP (10 μM)-induced TF protein (4 h) expressions in THP-1 cells were analyzed after the cells were pretreated with either ARC-118925XX (1µM) for 45min (**C**) or with Kaempferol (10µM) for 45min (**D**). Mouse survival proportions were determined after the wild-type, and P2Y2R-null mice were challenged with LPS (900 µg) via tail vein injections, after which mice death was observed for a total of five days. n=10 in each group (**E**). TF protein expression was determined by Western blotting assay in bone marrow cell lysates prepared from P2Y2R^+/+^ and P2Y2R^−/−^ mice after LPS (900 µg) challenge for 24 h. The data shown are representative of two independent experiments using bone marrow cells isolated from four mice in each group (**F**).

## Discussion

In the present study, we show for the first time that extracellular nucleotides, including UTP and ATP, stimulate TF expression both in protein and mRNA levels in human monocytic THP-1 cells and primary monocytes. We also show that ERK1/2, JNK, p38, and AKT pathways are all involved in nucleotides’ induction of TF expression. In addition, we have demonstrated that the P2Y2R is responsible for nucleotides signaling and TF induction in monocytes. Furthermore, we provided evidence that deletion of P2Y2R protects mice from endotoxemia death associated with decreased TF expression.

Despite the extensive prior studies focused on platelet P2Y12 receptor and thrombosis ^24–28^, research about the possible role of non-platelet P2Y receptors in the initiation and propagation of thrombosis are still lacking. However, accumulating evidence indicates the pathological roles of non-platelet P2Y receptors in the pathogenesis of inflammation *in vitro* and *in vivo* ^9,13,29–32^. This is particularly true for the P2Y2R. Under inflammatory conditions, P2Y2R can stimulate various downstream signaling pathways that regulate cell chemotaxis, proliferation, and migration in multiple types of cells ^33–38^. P2Y2R was shown to promote the expression of VCAM-1, increasing the adherence of monocytic cells to human endothelial cells ^15,39^. It is also known that P2Y2R promotes oxLDL-induced endothelial cell generation of reactive oxygen species and monocyte migration ^34^, suggesting that P2Y2R is involved in inducing intimal hyperplasia and monocyte infiltration in lesion formation in atherosclerosis. Our previous works demonstrated that ATP and UTP could induce TF upregulation through P2Y2R activation in human coronary artery endothelial cells ^19^. However, others have shown that monocyte TF is more critical than endothelial TF in provoking a pro-thrombotic state ^6,7^. Like endothelial cells, monocytes and macrophages have very little to no basal expression of TF in physiological conditions. However, TF expression can be induced by diverse inflammatory stimuli in monocytes/macrophages such as C-reactive protein, CD40 ligand, oxidized LDL, PDGF-BB, and endotoxin (LPS) ^3,6,7,40^. The present study shows that various nucleotides, including ATP, ADP, UTP, and UDP, can also induce TF expression in human monocytes through P2Y2R. It is conceivable that when significant monocyte surface TF is exposed and bound to plasma protein Factor VIIa, a pro-coagulable state of blood would be generated ^3,4,41^. Therefore, identifying novel factors regulating TF expression on blood cells might provide new insights into preventing pathological thrombosis. In addition to endothelial cells and monocytes/macrophages, other cells, such as smooth muscles cells, are alternatives that express inducible TF, which trigger increased cellular pro-coagulant activity ^3–5^. Although it is reported that the P2Y2R mediates coronary artery smooth muscle proliferation ^42^, it remains to be determined if smooth muscle TF could be upregulated by P2Y2R activation.

Several studies have highlighted the fundamental roles of P2Y receptors during infectious diseases over the past decade. Increasing evidence suggests that P2Y2R may be involved in LPS-mediated macrophage activation. LPS-stimulated monocytes and macrophages express elevated levels of P2Y2R mRNA ^13^, and P2Y2R activation increases neutrophil migration and IL-8 release from human monocytic THP-1 cells and U937 cells ^43^. LPS induces TF expression ^21^ and causes ATP release in THP-1 cells ^13^. However, whether ATP release and P2Y2R activation contribute to LPS-induced TF induction has not been studied. Surprisingly, the present study demonstrated that the P2Y2R appeared to play an unexpected “permissive” role for LPS-induced TF expression. Several lines of evidence support this notion: First, we found that removing extracellular nucleotides by apyrase largely inhibited LPS-induced TF induction; Second, blocking the P2Y2R by two structurally different P2Y2R-selective antagonists ARC-118925XX and Kaempferol similarly abolished LPS-induced TF upregulation; Third, siRNA silencing P2Y2R completely suppressed LPS-induced TF expression; Finally, challenging the P2Y2R-null mice with LPS in vivo produced no to a negligible amount of TF expression in bone marrow cells as compared with LPS action on the wild-type mice. Thus, we provided compelling evidence to support a vital role of P2Y2R not only in nucleotides-induced but also in LPS-induced TF upregulation in monocytes. Consistent with these findings, we found that P2Y2R-deficient mice had a greater survival rate than wild-type mice in response to a lethal dose of LPS challenge. Proctor et al. discovered that 2-Methylthio-ATP ^18^, a ligand for multiple P2X and P2Y receptors, reduces LPS-stimulated TNF-α and IL-1β release *in vivo* and protects mice from LPS-induced death, implying that the coordinated action of P2Y and P2X receptors are essential for modulating LPS responses *in vivo* ^13^. However, these earlier studies have not determined the involved subtype(s) of P2X or P2Y receptors. In addition, it remains unknown if the pharmacological effect of 2-Methylthio-ATP they reported was due to its agonist or antagonistic action. Our preliminary data indicate that 2-Methylthio-ATP indeed acted as a weak agonist on P2Y2R and blocked ATP activation of P2Y2R (data not shown).

Prior studies indicated that extracellular nucleotides regulate Toll-like receptor functions such as TLR4 signaling in human endothelial progenitor cells ^44^. Consistent with the previous study, we show that LPS stimulates a dramatic release of ATP in THP-1 cells. Several studies revealed that some other inflammatory stimuli, such as oxLDL, can cause a rapid release of ATP from human endothelial cells ^34^, and that a high concentration of extracellular ATP itself could induce ATP release with a potential mechanism leading to amplification of ATP signaling ^45^. Therefore, it is conceivable that P2Y2R activation by released and accumulated ATP/UTP due to inflammation and/or cell death may be a significant component of total TF induction triggered by many stimuli (a working model presented online Fig. 2). In this perspective, identifying a novel role for the P2Y2R in monocyte TF induction may have unique importance in advancing our knowledge on pathological thrombosis, including but not limited to disseminated intravascular coagulation.

The MAPK pathways are implicated in TF induction in several systems, but each member of the MAPK family contributes to TF induction in an extracellular stimulus-and cell type-dependent manner ^20,21^. However, in the current study, we found that all the p38, JNK, ERK1/2, and AKT pathways are involved in P2Y2R-induced TF upregulation in THP-1 cells. These findings are inconsistent with what we reported in human coronary artery endothelial cells in which we did not find a positive role of ERK1/2 or AKT in P2Y2R-induced TF expression ^19,20^. Indeed, we found that ERK1/2 plays a negative role through the transcription factor Fra-1 in human endothelial cells ^46^, and AKT is not activated at all by endothelial P2Y2R ^19^. The exact mechanism(s) responsible for this discrepancy is unknown. However, in addition to different cell backgrounds, it should be noted that in human coronary artery endothelial cells, we did not detect any c-Fos expression and its phosphorylation in response to P2Y2R activation ^20^. In contrast, in monocytes, we found that stimulation of the P2Y2R activated not only Fra-1, ATF-2, and c-Jun, but also c-Fos. Nonetheless, it remains to be determined how these four transcriptional factors interplay to control P2Y2R-induced TF upregulation in monocytes. On the other hand, the PI3K/AKT pathway was reported to negatively regulate endothelial TF expression as inhibition of PI3K, or its downstream mediators increased TF expression in response to TNF-α, histamine, thrombin, and VEGF ^4^. Here, we observed that AKT is actively involved in P2Y2R-mediated monocyte TF induction; this is not unexpected since others found that the PI3K/AKT pathway mediates TF expression in MDA-MB-231 breast cancer cells ^47^. Thus, it appears that the AKT pathway plays a cell-type dependent role in the control of TF expression.

In addition to exploring cell signaling mechanisms, we also discovered that ATP and UTP upregulate the TF pre-mRNA and mature mRNA expression in THP-1 cells, suggesting a post-P2Y2R transcriptional mechanism for TF induction. However, it should be noted that post-transcriptional mechanisms were postulated in control of TF expression; indeed, LPS induces TF up-regulation via increased TF mRNA stability ^21^. The exact molecular mechanism(s) responsible for increased TF mRNA stability remains unknown. In line with this, we could not exclude the possibility that P2Y2R-mediated TF mRNA up-regulation in monocytes may be partially due to mRNA stabilization. We wish to dissect the transcriptional and post-transcriptional components in a follow-up study.

In summary, we report the first evidence that activating the P2Y2R by various extracellular nucleotides induces TF expression in monocytes and that the MAPK and AKT pathways contribute to P2Y2R-induced TF expression. In addition, P2Y2R deficiency protects the mice from endotoxemia death associated with a decreased TF expression in bone marrow cells. Our finding suggests that in addition to the platelet P2Y12 receptor, non-platelet P2Y receptor(s), e.g., the monocyte P2Y2R, is a potential new drug target in the prevention and/or treatment of various TF-related diseases such as sepsis, atherosclerosis, and cancers.

## Acknowledgments

The content is solely the authors’ responsibility and does not necessarily represent the official views of the National Institutes of Health.

---

**Supplemental Figure 1.**
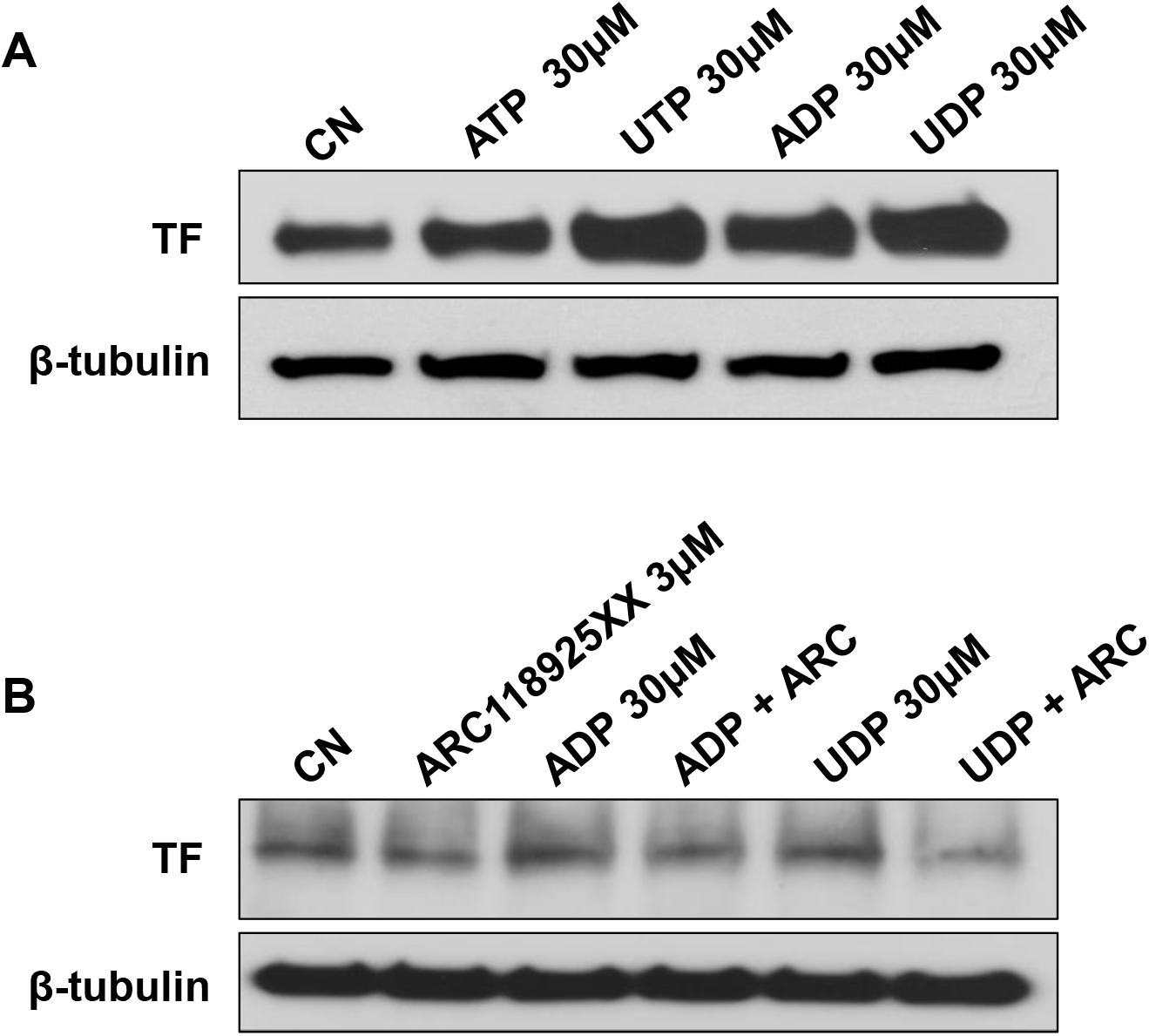
Role of P2Y2R in ADP/UDP-induced TF expression in THP-1 monocytes. The Western blotting assay analyzed ATP-, UTP-, ADP-, or UDP (30µM)-induced TF protein expressions in THP-1 cells after the indicated nucleotides stimulated the cells for 4 h (**A**). ADP-or UDP-induced TF protein expressions were analyzed after the cells were pretreated with or without ARC-118925XX (3µM) for 45min followed by ADP or UDP stimulation of the cells for 4 h. The data shown are representative of three independent experiments (**B**).

**Supplemental Figure 2.**
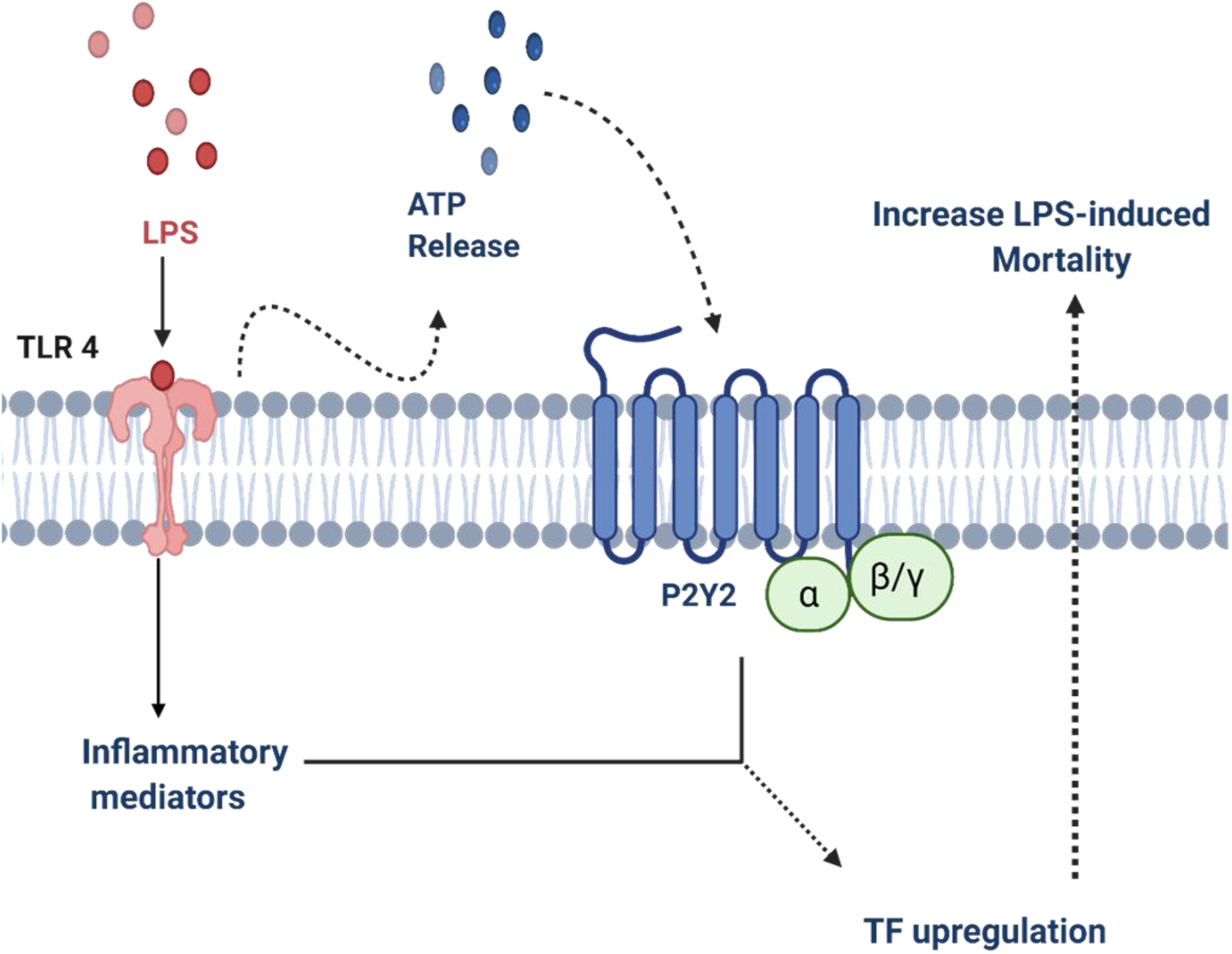
A proposed working model for P2Y2R mediation of LPS-induced TF upregulation and endotoxemia death. LPS-induced ATP release and P2Y2R activation in monocytes are indispensable components for LPS-induced monocyte TF upregulation and endotoxemia death in mice.

**Supplemental Table 1:**
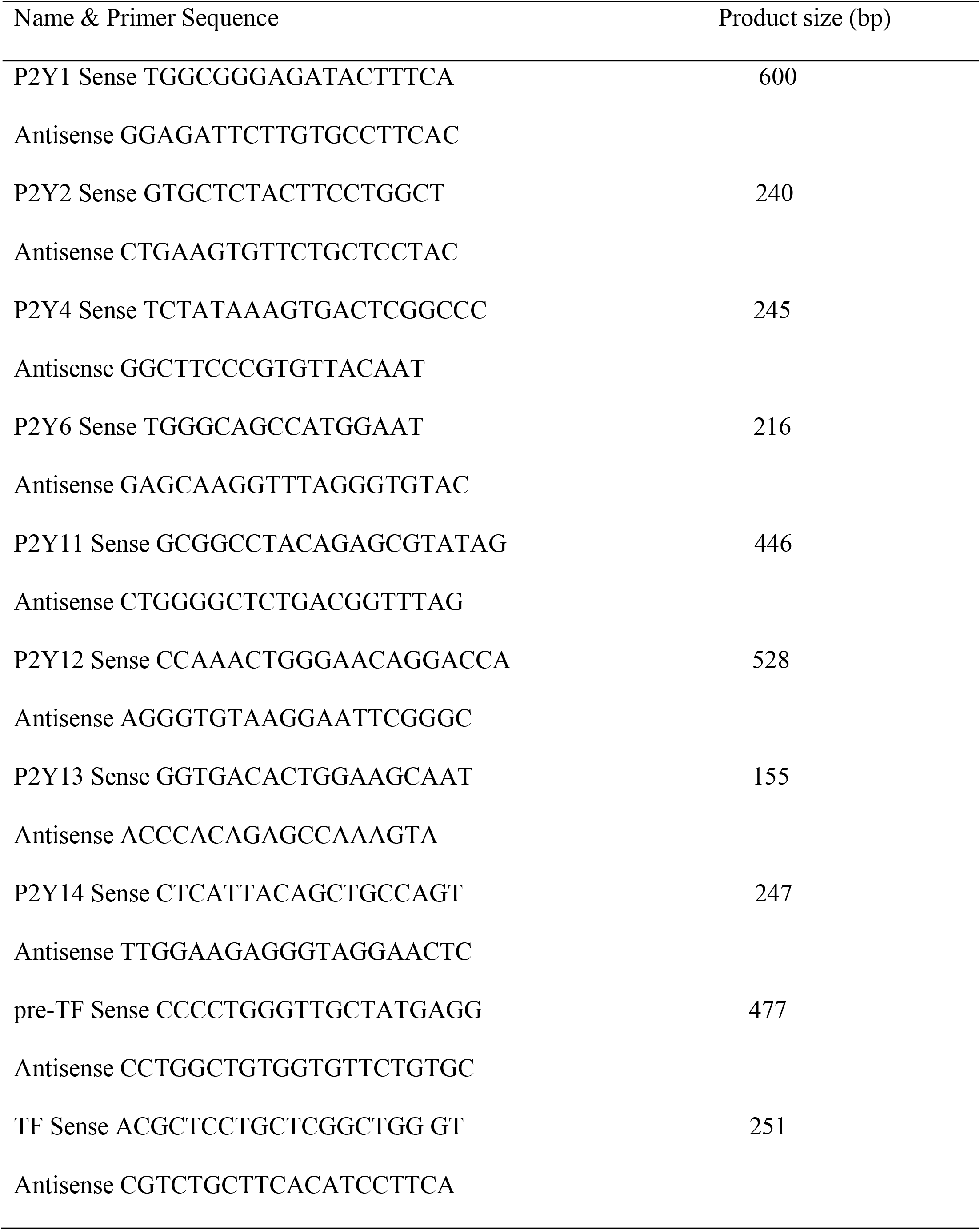
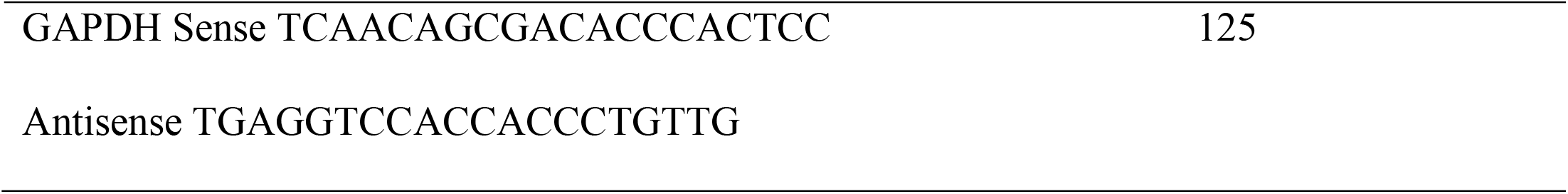
PCR primer sequences and their product sizes.

